# Time-Resolved SAXS Unveils Two Structurally Distinct Membranes upon Cytochrome c Binding

**DOI:** 10.64898/2026.11.13.765644

**Authors:** Leandro R. S. Barbosa, Alessio Romaldini, Alessio Correani, Andrea Travaglini, Paolo Mariani, Rosangela Itri, Francesco Spinozzi

## Abstract

The association of peripheral proteins with lipid membranes can couple lateral lipid organization to interbilayer correlations, but the temporal evolution of these structural changes remains difficult to characterize experimentally. Here, we use time-resolved small-angle X-ray scattering (SAXS) to investigate the structural evolution of POPC:POPG large unilamellar vesicles following the addition of cytochrome c. The experimental profiles are analyzed globally using an extension of the SASBIN model incorporating molecularly constrained bilayer electron-density profiles, lateral lipid inhomogeneities, and interbilayer correlations. A two-population description consistently reproduces the SAXS profiles, with distinct structural contributions associated with protein-free and cytochrome-c-associated bilayers. Within this framework, the protein-associated population exhibits lateral correlations consistent with POPG-rich domains and a progressive increase in lamellar ordering, whereas the protein-free population is characterized by molecular-scale effective inhomogeneities. The fitted time-dependent parameters suggest that population redistribution and lateral reorganization are most pronounced during the early stages of the interaction, while some parameters describing interbilayer correlations evolve more gradually. These results provide a quantitative structural description of how cytochrome c association can couple lateral lipid organization and lamellar ordering in mixed zwitterionic/anionic membranes.

## 1. Introduction

Cytochrome c (Fig. 1A) is a small soluble heme protein whose function extends well beyond its classical role as an electron carrier in the mitochondrial respiratory chain. Under physiological conditions, cytochrome c shuttles electrons between complexes III and IV in the mitochondrial intermembrane space. Under cellular stress, however, its redistribution and interaction with mitochondrial membranes become central events in the regulation of programmed cell death [1]. In particular, membrane-associated cytochrome c participates in the sequence of processes that links mitochondrial membrane remodeling, cardiolipin oxidation, mitochondrial outer membrane permeabilization, and release of cytochrome c into the cytosol [1, 2, 3, 4]. These processes are not controlled only by the intrinsic properties of the protein, but also by the composition, charge density, hydration, and lateral organization of the lipid bilayer. Cytochrome c binding to anionic phospholipids therefore represents a paradigmatic example of how peripheral proteins can couple electrostatic recognition, local membrane remodeling, lipid redistribution, and biochemical activation.

**Figure 1:**
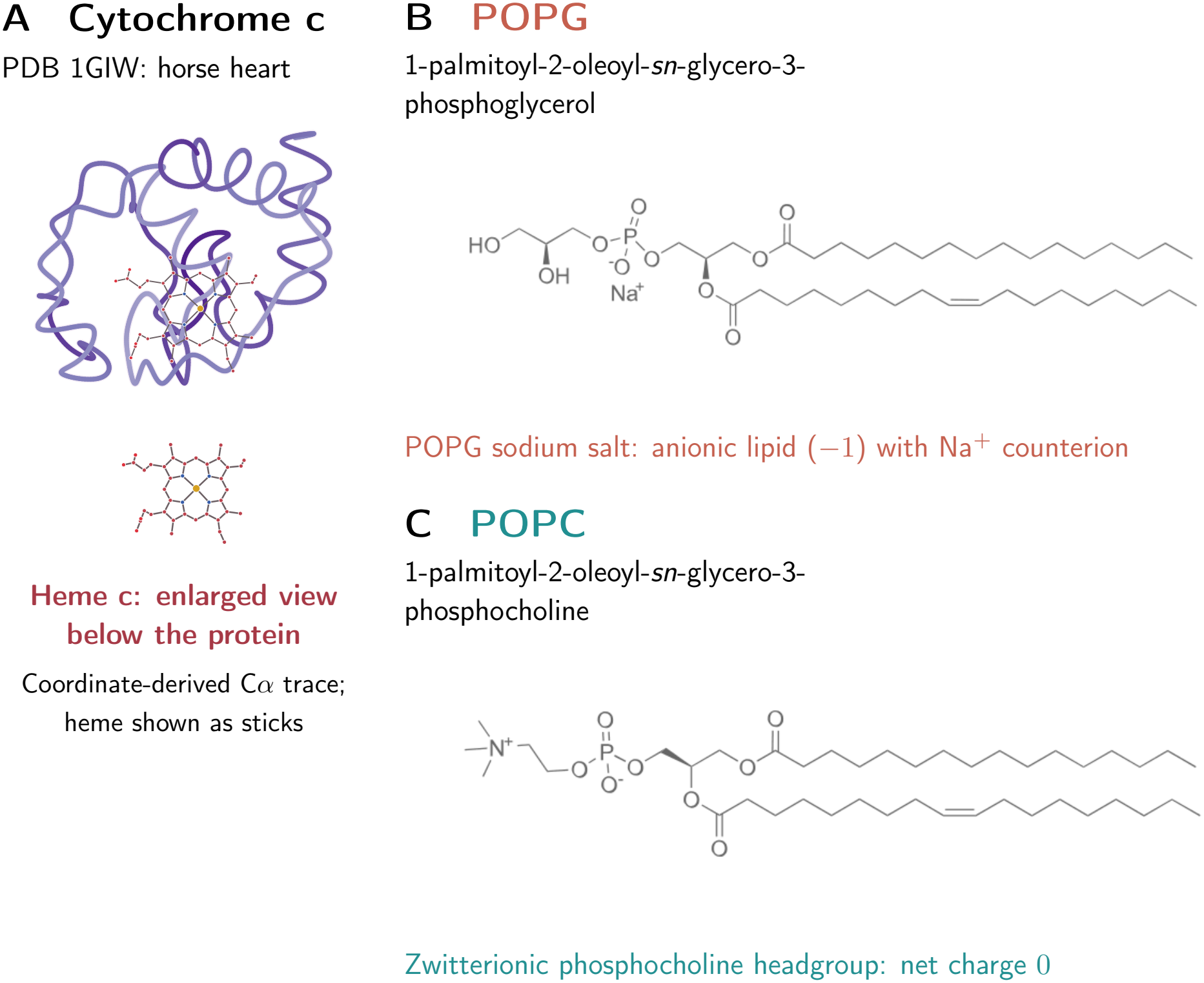
Molecular structures of the components used in this study. (A) Horse-heart cytochrome c represented by a smoothed C*α* trace derived from PDB 1GIW, with the heme c group shown in ball-and-stick representation and enlarged below. (B) POPG sodium salt and (C) zwitterionic POPC, reproduced from the author-supplied chemical structures. Both phospholipids contain a palmitoyl chain (16:0) at the *sn*-1 position and an oleoyl chain (18:1, cis-Δ^9^) at *sn*-2. Molecular representations are not to a common scale.

Among anionic phospholipids, cardiolipin has received particular attention because of its high abundance in the mitochondrial inner membrane and its specific involvement in mitochondrial function, immune signaling, and cell death [2, 4]. Cardiolipin is a tetra-acylated diphosphatidylglycerol whose oxidation and redistribution across mitochondrial membranes are closely associated with apoptotic signaling [2, 3, 4]. The interaction between cytochrome c and cardiolipin has been proposed to involve electrostatic contacts between positively charged protein patches and negatively charged lipid headgroups. This interaction can convert cytochrome c from an electron-transfer protein into a lipid peroxidase, promoting cardiolipin oxidation during the early stages of apoptosis [3, 5]. More generally, oxidized phospholipids are bioactive lipid species capable of modifying membrane physical chemistry properties and protein recognition processes [3, 6]. These observations highlight the importance of understanding not only whether cytochrome c binds to anionic membranes, but also how its binding modifies the structural organization of the membrane itself.

Because mitochondrial membranes are chemically and structurally complex, simplified model systems are essential for isolating the mechanisms that control cytochrome c-membrane interactions. Phosphatidylcholine (PC) lipids are zwitterionic phospholipids and constitute one of the most common matrix components used in biomimetic membranes. In contrast, phosphatidylglycerol (PG) lipids carry a negatively charged headgroup and therefore provide a convenient model for the electrostatic component of cytochrome c binding to anionic membranes. In the lipid abbreviations used here, PO denotes the palmitoyl-oleoyl acyl-chain composition, i.e. one saturated palmitoyl chain and one monounsaturated oleoyl chain, a common mixed-chain composition in fluid phospholipid bilayers. Accordingly, 1-palmitoyl-2-oleoyl-sn-glycero-3-phosphocholine (POPC, Fig. 1C) provides a zwitterionic fluid bilayer matrix, whereas 1-palmitoyl-2-oleoyl-sn-glycero-3-phosphoglycerol (POPG, 1B) introduces a controlled fraction of anionic lipid into the same hydrocarbon tail.

POPC:POPG mixtures are therefore well suited to investigate how a zwitterionic membrane matrix responds to the incorporation of an anionic lipid species and to the binding of a positively charged peripheral protein. The use of lipids sharing the same palmitoyl-oleoyl chains minimizes differences associated with hydrophobic-chain packing, thereby focusing the comparison on headgroup charge, hydration, ion binding, and lateral lipid organization. Recent studies of cytochrome c in anionic phospholipid nanodiscs have further supported the use of POPG as a model lipid for probing cytochrome-c-membrane interactions [7]. Nevertheless, comparisons between PG and cardiolipin show that the structural and functional response of cytochrome c can depend on the specific anionic lipid, with different extents of protein destabilization and peroxidase activation [5].

This point is particularly relevant because charged lipid membranes can display complex structural behavior. Ionic conditions can affect headgroup hydration, cation binding, membrane compression, and lipid packing, even in zwitterionic PC bilayers [8]. Anionic phosphatidylglycerol membranes can show even richer behavior, including ion-and concentration-dependent phase behavior, lateral organization, membrane perforation, and domain formation [9]. Consequently, even apparently simple POPC:POPG mixtures may exhibit lateral heterogeneity whose evolution is sensitive to protein binding. A key unresolved question is therefore whether cytochrome c binds to a pre-existing lateral organization, selectively stabilizes anionic lipid-rich regions, or actively promotes lipid redistribution and the emergence of distinct membrane populations.

To address this issue, we make use of Small-angle X-ray (SAXS) that is a powerful tool for probing lipid bilayers at nanometric length scales. SAXS is particularly sensitive to the electron-density contrast between the polar headgroup region, the hydrocarbon-chain region, and the terminal methyl region [10, 11, 12]. However, the structural interpretation of scattering data from fully hydrated fluid bilayers is intrinsically model dependent. Fluid bilayers are not crystalline objects with sharply defined atomic positions, but fluctuating structures described by broad statistical distributions. Therefore, physically meaningful parameters such as bilayer thickness, area per lipid, hydration, and component electron densities must be inferred through models constrained by molecular volumes, chemical composition, and known structural relationships [10, 11, 13, 14]. This requirement becomes even more stringent when the membrane contains more than one lipid species or when lateral inhomogeneities are present.

Several levels of structural modeling have been developed to analyse lipid bilayers by small-angle scattering. Early and widely used descriptions treated the membrane as a onedimensional electron-density or scattering-length-density profile, often represented by a small number of slabs with uniform density. In its simplest form, this description separates the bilayer into polar headgroup regions, hydrocarbon-chain regions, and a terminal-methyl region, each characterized by its own thickness and density. Such slab models provide a transparent connection between the measured form factor and the average bilayer profile, but they require physical constraints to avoid non-unique or unphysical solutions [14, 15]. More refined approaches, such as the scattering density profile model and its extensions, introduced molecular volume constraints and quasi-molecular fragments, allowing bilayer structural parameters to be extracted from SAXS with improved physical chemistry consistency [10, 11, 12]. Along the same line, the modified scattering density profile approach provided a smooth molecularly resolved description of oxidized POPC membranes, enabling SAXS data to be interpreted in terms of the location of specific chemical groups and oxidation-induced changes in area per lipid and membrane thickness [16]. These approaches emphasized that quantities such as area per lipid, hydration, and bilayer thickness cannot be determined independently from scattering data alone, but are linked through molecular volumes, chemical composition, and conservation relationships.

Despite these advances, conventional one-dimensional bilayer models remain laterally averaged. They are therefore not designed to distinguish whether changes in the scattering curve arise from homogeneous modification of the average bilayer profile or from the formation of laterally organized regions with different electron density. This limitation is central for mixed lipid membranes interacting with peripheral proteins, where binding may recruit specific lipids, stabilize local domains, alter headgroup hydration, and generate coexisting membrane populations. In such systems, a model that explicitly accounts for in-plane scattering-length-density inhomogeneities is required.

The SASBIN model was developed to address this problem by representing scattering-length-density inhomogeneities, such as lipid domains, pores, and membrane-associated proteins, as laterally correlated cylindrical objects embedded in a host bilayer, with the possibility of including lamellar stacking [17]. In SASBIN, the bilayer and the inhomogeneities are described by slab-like electron-density profiles in which the transitions between adjacent levels are smoothed, avoiding sharp discontinuities while preserving an analytically tractable description of the form factor.

In the current work, we apply time-resolved SAXS to investigate the structural evolution of POPC:POPG large unilamellar vesicles upon addition of cytochrome c. The data are analysed with the SASBIN model, implemented in GENFIT [17, 18], and extended here by imposing a molecular description of the bilayer electron-density profile. In this description, the thicknesses and electron densities of the polar, hydrocarbon-chain, and terminal-methyl regions are calculated from molecular composition, group volumes, electron counts, hydration, and area per lipid, rather than treated as independent phenomenological parameters. The model also allows for the coexistence of distinct bilayer populations, lateral POPG-rich inhomogeneities, lamellar stacking, and membrane-associated cytochrome c.

The central question is how cytochrome c association couples lateral membrane organization to interbilayer correlations in a mixed zwitterionic/anionic lipid membrane. POPC:POPG vesicles provide a controlled model system for investigating these processes, without attempting to reproduce the full lipid composition of mitochondrial membranes.

## 2. Material and Methods

### 2.1. Sample preparation

POPC (1-palmitoyl-2-oleoyl-*sn*-glycero-3-phosphocholine) and POPG (1-palmitoyl-2-oleoyl-*sn*-glycero-3-phosphoglycerol) were purchased from Avanti Polar Lipids (Alabaster, AL, USA) and used without further purification. Large unilamellar vesicles (LUVs) were prepared from an equimolar POPC:POPG mixture (1:1 molar ratio). The appropriate amounts of each lipid were dissolved in chloroform in a round-bottom flask, and the organic solvent was removed under a gentle stream of ultrapure nitrogen gas to produce a homogeneous thin lipid film on the flask wall. The resulting film was subsequently kept under low pressure overnight to ensure removal of residual chloroform.

The dried lipid film was hydrated with 20 mM phosphate buffer at pH 7.0 to form a lipid dispersion. After hydration, the suspension underwent repeated freeze-thaw cycles (*≈* 6), with thawing performed above the phase-transition temperatures of the constituent lipids to promote efficient membrane reorganization and improve vesicle homogeneity. The resulting multilamellar lipid suspension was then converted into LUVs by sequential extrusion through polycarbonate membranes (Avanti Polar Lipids, Alabaster, Alabama, USA). Each sample was first passed 31 times through a membrane with a nominal pore diameter of 400 nm and subsequently subjected to an additional 31 extrusion steps through a 100 nm membrane. This procedure yielded vesicles with a nominal diameter of approximately 100 nm.

Cytochrome c from equine heart (C7752, Merck KGaA, Darmstadt, Germany) was used as received, without additional purification or any externally imposed oxidation or reduction treatment. Protein stock solutions were prepared in the same 20 mM phosphate buffer at pH 7.0 used for lipid dispersions. For protein–membrane interaction experiments, LUV suspension and cytochrome c solution were combined in a 1:1 volume ratio, typically by mixing approximately 0.5 mL of each solution, resulting in a final sample volume of approximately 1.0 mL.

After mixing, the final total lipid concentration was 7 mM, and the final cytochrome c concentration was 10*mg/mL*. The protein concentration was independently verified by spectrophotometry. Using an approximate molecular mass of 12 kDa for equine heart cytochrome c, 10 mg/mL corresponds to approximately 0.83 mM protein. Unless otherwise specified, sample preparation and all subsequent measurements were performed at room temperature.

### 2.2. SAXS experiments

Small-angle X-ray scattering (SAXS) experiments were performed at the SAXS-1 beam-line of the Brazilian Synchrotron Light Laboratory (LNLS/CNPEM), Campinas, Brazil. Measurements were conducted at room temperature using monochromatic X-rays with a wavelength of 1.488 Å. The scattered intensity was recorded using a Pilatus 300K two-dimensional detector (Dectris), positioned approximately 1000 mm from the sample. The experimental configuration provided access to a scattering vector modulus range of approximately 0.01–0.30 Å^*−*1^, where *q* = (4*π/λ*) sin *θ, λ* is the X-ray wavelength, and 2*θ* is the scattering angle. Approximately 300 *µ*L of each sample was introduced into a sample holder equipped with mica windows. SAXS measurements of POPC:POPG (1:1 molar ratio) lipid vesicles were initially performed in the absence of cytochrome c to establish the structural characteristics of the lipid membranes before protein addition. For the protein–membrane interaction experiments, the vesicle suspension and cytochrome c solution were manually combined at a 1:1 volume ratio, resulting in final concentrations of 7 mM total lipid and 10 mg/mL cytochrome c.

Immediately after mixing, an aliquot of the resulting suspension was transferred to the SAXS sample holder and positioned in the X-ray beam. The sample remained in the same holder throughout the entire time-dependent experiment, without replacement, further mixing, or additional manipulation between consecutive measurements. This experimental arrangement allowed the structural evolution of the same protein–lipid dispersion to be monitored continuously over the incubation period through a sequence of discrete SAXS acquisitions, thereby avoiding sample-to-sample variations that could arise from repeated preparation or replacement. Time-dependent SAXS measurements were performed using a repeated acquisition protocol consisting of an X-ray exposure of approximately 40 s, followed by a waiting interval of approximately 5 min before the next exposure. During these waiting periods, no scattering data were collected, and the sample remained stationary in the sample holder under the same experimental conditions. This acquisition–waiting sequence was repeated throughout the experiment to investigate the progressive structural changes induced by the interaction of cytochrome c with the POPC:POPG lipid bilayers. Each recorded SAXS profile therefore represents the scattering intensity integrated over a 40 s acquisition window, rather than an instantaneous measurement of the membrane structure. Because sample mixing, transfer, positioning, and initiation of the first exposure were performed manually, a finite experimental dead time was present between the initial contact of the protein and lipid dispersions and the acquisition of the first SAXS profile. This interval was estimated to be approximately 60 *±* 20 s. Although the instant of mixing represents the physical onset of the protein–membrane interaction, this initial delay was not explicitly included in the nominal experimental times used for the analysis of the SAXS data.

Consequently, the reported incubation times correspond to the nominal acquisition schedule, while the actual interaction time includes an additional initial offset of approximately 60–80 s. The temporal sampling interval was therefore determined by the combination of the 40 s exposure duration and the approximately 5 min waiting period between successive acquisitions. Structural rearrangements occurring during the initial experimental dead time could not be independently resolved, and changes taking place within an individual exposure were averaged in the corresponding scattering profile. Similarly, structural transformations occurring between consecutive measurements were monitored only through their effects on the subsequent SAXS patterns. Nevertheless, this acquisition protocol provided a consistent means of following the longer-timescale evolution of lipid membrane organization following cytochrome c addition while maintaining identical sample conditions throughout each experimental series. All SAXS experiments were carried out at room temperature, and the same experimental geometry and acquisition protocol were maintained for the corresponding time-dependent measurements. Prior to the time-dependent SAXS experiments, control measurements were performed on cytochrome c solutions in the absence of lipids and on POPC:POPG vesicle suspensions in the absence of protein. All samples, including the corresponding protein–lipid mixtures, were systematically evaluated for potential radiation-induced damage by monitoring changes in their SAXS profiles upon X-ray exposure. No detectable radiation-induced alterations were observed under the experimental conditions employed, consistent with the relatively low incidence of radiation damage typically encountered under comparable measurement conditions at the Brazilian synchrotron facility.

### 2.3. SAXS Data Analysis

The SAXS data were analyzed using the SASBIN model implemented in the GENFIT software package [17, 18]. The model describes the experimental scattering curves by assuming the coexistence of two structurally distinct POPC:POPG bilayer populations. Population 1 consists of bilayers without membrane-associated cytochrome c, whereas population 2 contains cytochrome c molecules confined between adjacent bilayers. The fraction of bilayers belonging to population 1 is denoted by *x*_1_.

Within the SASBIN formalism, all lateral inhomogeneities are represented by cylindrical objects spanning the entire thickness of the corresponding scattering entity and composed of regions with uniform electron density. In the present work, the lipid inhomogeneities are identified with cylindrical POPG-rich domains embedded in a POPC-rich matrix, whereas cytochrome c is represented by an equivalent core-shell cylinder. This common geometrical description provides a unified framework for calculating the scattering contributions arising from both lipid domains and membrane-associated proteins.

The structural model adopted for the two bilayer populations is schematically illustrated in Fig. 2A-D. In both populations, the lipid bilayer is described by a symmetric three-level electron-density profile, for each leaflet consisting of the hydrated polar headgroup region (*i* = 1), the hydrocarbon-chain region (*i* = 2), and the terminal methyl region (*i* = 3).

**Figure 2:**
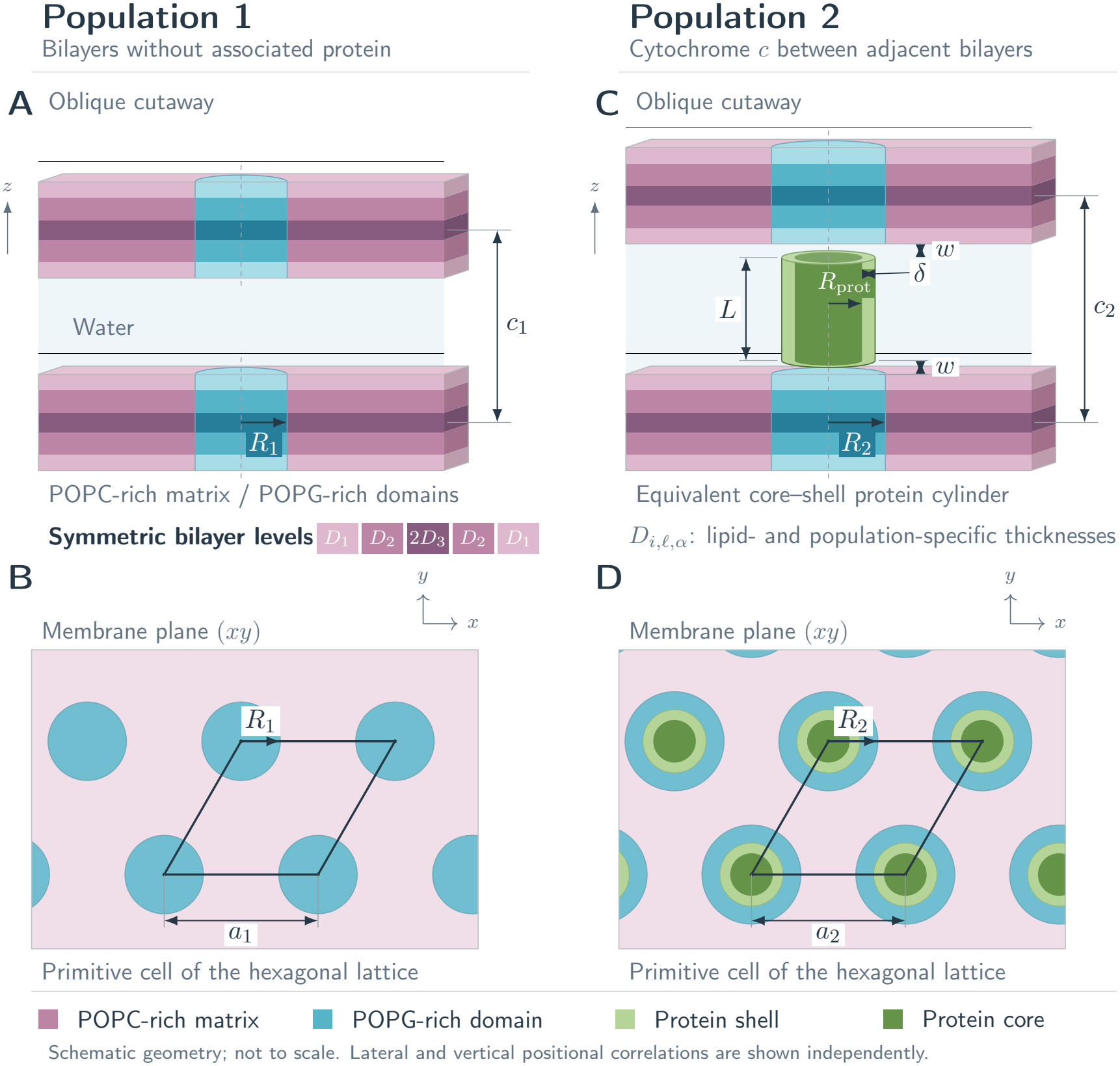
Schematic representation of the two bilayer populations in the SASBIN analysis. (A,C) Oblique cutaway views illustrating vertical correlations in populations 1 and 2, respectively. (B,D) Corresponding projections onto the membrane plane, with cylindrical POPG-rich domains embedded in a POPC-rich matrix. The outlined rhombus is a primitive cell of the hexagonal lattice, whose parameter is *a*_*α*_; domain radii are denoted by *R*_*α*_. Each bilayer has three symmetric electron-density levels, arranged as *D*_1_, *D*_2_, 2*D*_3_, *D*_2_, *D*_1_. Their thicknesses and densities are lipid-and population-specific; the colours identify lipid regions rather than numerical density values. Population 2 contains an equivalent core–shell cytochrome *c* modeled as a cylinder of height *L*, core radius *R*_prot_, and radial shell thickness *δ*. Water layers of thickness *w* separate the protein from the adjacent POPG-rich domains. The bilayer midplane spacings are *c*_1_ and *c*_2_, with 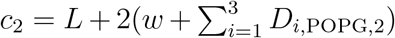. Idealized correlations are shown schematically; the drawing is not to scale.

In the present implementation, both lipid species were allowed to adopt population-specific molecular structures. Thus, POPC and POPG were each described by two sets of structural parameters, one associated with the protein-free population and one associated with the cytochrome-c-associated population. This choice accounts for the fact that cytochrome-c binding and the formation of POPG-rich domains may also modify the POPC-rich matrix surrounding the anionic domains.

For each population *α* = 1, 2, the electron-density profiles of the POPC-rich matrix and of the POPG-rich domains were therefore constructed from population-specific molecular parameters. The corresponding thicknesses and electron densities of the three bilayer regions were calculated separately for POPC_*α*_ and POPG_*α*_ from the molecular areas, hydration numbers, group volumes, and numbers of electrons defined below.

The principal extension of the original SASBIN model introduced in the present work concerns the molecular description of the bilayer electron-density profile. Instead of refining independently the thicknesses and electron densities of the three electron-density levels, these quantities are calculated from the molecular composition of the lipid molecules. The molecular volumes (*ν*_*j*_), numbers of electrons (*e*_*j*_), and multiplicities (*N*_*j,l*_) of the chemical groups employed in the present model are summarized in Table 1, where *l* = POPC or POPG denotes the lipid species. The multiplicity *N*_*j,l*_ represents the number of occurrences of the *j*th chemical group in one molecule of lipid *l*. The hydrated polar headgroup region (domain 1) contains all polar chemical groups together with the associated hydration water. Since both lipid species are allowed to adopt population-specific molecular structures, the molecular volumes and numbers of electrons were calculated separately for each lipid species *l* = POPC, POPG and for each bilayer population *α* = 1, 2. Accordingly,

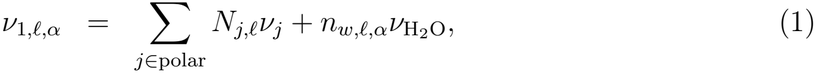

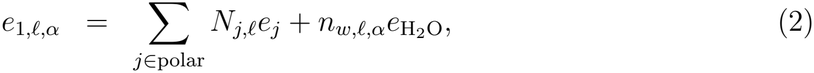

where *n*_*w,l,α*_ is the number of water molecules hydrating the headgroup of lipid species *l* in population *α*. Here, 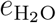 and 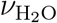 denote the number of electrons and the molecular volume of a water molecule, respectively (Table 1).

**Table 1:** Molecular volumes (*ν*_*j*_), numbers of electrons (*e*_*j*_), and multiplicities (*N*_*j,l*_) of the chemical groups used to construct the electron-density profiles of POPC and POPG, where *l* denotes the lipid species. The hydrophobic chains are described in terms of CH, CH_2_, and CH_3_ groups. The polar headgroup is represented by quasi-molecular fragments. CG denotes the carbonyl-glycerol fragment (CH_2_)(CH)(COO)(CH_2_)(COO); PCN denotes the fragment (PO_4_)(CH_2_)(CH_2_)N, comprising the phosphate group, the two methylene groups linking the phosphate to choline, and the quaternary nitrogen; CholCH_3_ represents the three methyl groups attached to the quaternary nitrogen; PGH denotes the phosphoglycerol moiety (PO_4_)(CH_2_)(CH)(OH)(CH_2_)(OH) of the POPG headgroup.

| Chemical group | *ν*<sub>*j*</sub> | *e*<sub>*j*</sub> | *N*<sub>*j*,*ℓ*</sub> |  |
|---|---|---|---|---|
|  | (Å<sup>3</sup>) |  | POPC | POPG |
| *Hydrophobic groups* |  |  |  |  |
| CH | 21.5 | 7 | 2 | 2 |
| CH<sub>2</sub> | 27.7 | 8 | 28 | 28 |
| CH<sub>3</sub> | 52.9 | 9 | 2 | 2 |
| *Polar groups* |  |  |  |  |
| CG | 143.3 | 67 | 1 | 1 |
| PCN | 87.8 | 70 | 1 | 0 |
| CholCH<sub>3</sub> | 99.9 | 27 | 1 | 0 |
| PGH | 147.7 | 88 | 0 | 1 |
| *Water* |  |  |  |  |
| H<sub>2</sub>O | 29.9 | 10 |  |  |

Because of the conformational disorder of the hydrocarbon chains, the terminal methyl groups are not confined to the bilayer midplane, whereas part of the methylene and olefinic CH groups penetrates into the terminal methyl region. Consequently, the hydrophobic core is described by two overlapping electron-density levels rather than by chemically distinct methylene and terminal methyl layers. For each lipid species *l* = POPC, POPG and for each bilayer population *α* = 1, 2, the molecular volumes associated with domains 2 and 3 are therefore

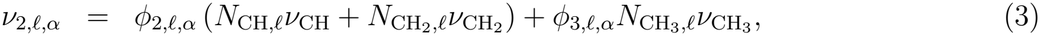

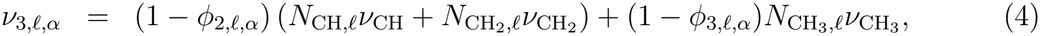

whereas the corresponding numbers of electrons are

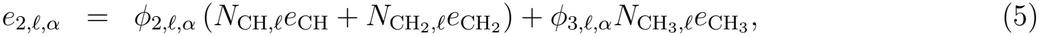

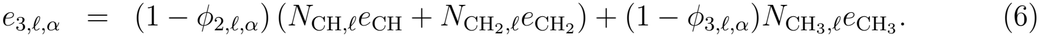

The electron densities of the three electron-density levels are then

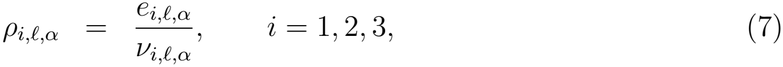

whereas the corresponding thicknesses are calculated from the molecular area per lipid head-group,

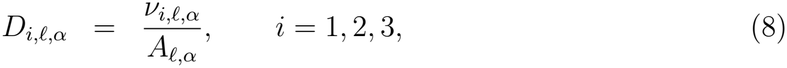

where *A*_*l,α*_ denotes the molecular area per lipid headgroup of lipid species *l* in population *α*. In population 2, cytochrome c is assumed to be preferentially associated with the POPG-rich domains because of the electrostatic attraction between the negatively charged PG headgroups (net charge *−*1 per lipid at neutral pH) and the positively charged protein, whose net charge is approximately +10 elementary charges according to its primary amino-acid sequence. The parameter *w* represents the thickness of the water layer separating the protein from the POPG-rich domain (Fig. 2C).

Within the SASBIN formalism, all scattering inhomogeneities are represented by geometrical objects composed of regions with uniform electron density. For consistency with this description, the form factor of cytochrome c was not directly calculated from its atomic coordinates during the fitting procedure. Instead, the atomic coordinates of PDB entry 1GIW were first analyzed with the SASMOL method [19] to obtain the corresponding scattering profile. This profile was then represented by an equivalent core-shell cylinder, following the parameterization reported in Table 3 of Spinozzi et al. [17]. The geometrical parameters of the cylinder, namely the core radius *R*_prot_, the shell thickness *δ*, and the cylinder height *L*, together with the volume fractions *ϕ*_*i*_ and *ϕ*_*e*_ of polypeptide matter in the core and shell regions, respectively, were fixed to the values obtained from this parameterization and were not refined during the fitting procedure.

For each bilayer population, the membrane organization is described in terms of independent lateral and vertical correlations. The lateral organization of the cylindrical POPG-rich domains is described by the hexagonal lattice parameter *a*_*α*_ and the paracrystalline distortion parameter *g*_*a,α*_, where *α* = 1, 2 identifies the two bilayer populations.

The vertical organization was treated by allowing, within each bilayer population, the coexistence of vertically correlated and vertically uncorrelated bilayers. Accordingly, for each population *α*, two SASBIN contributions were calculated. The first one, denoted by 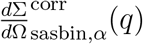, was calculated using the full vertically correlated SASBIN expression, with an average number of correlated bilayers *N*_*c,α*_ *>* 1. The second one, denoted by 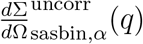, was calculated by setting *N*_*c,α*_ = 1, so that the corresponding vertical structure factor reduces to unity. The correlated and uncorrelated contributions of a given population share the same molecular bilayer profile and the same lateral organization parameters; they differ only in the vertical structure factor.

The vertical organization of the correlated component of population *α* is characterized by the correlation distance *c*_*α*_, the average number of correlated bilayers *N*_*c,α*_, and the paracrystalline distortion parameter *g*_*c,α*_. For population 1, *c*_1_ is an independent fitting parameter. In population 2, the interlamellar distance is determined by the geometry of the protein-mediated bilayer stack and is therefore given by.

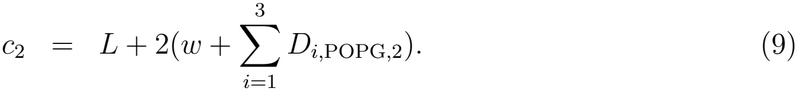

For the uncorrelated contributions, *N*_*c,α*_ is fixed to unity; therefore, the corresponding values of *c*_*α*_ and *g*_*c,α*_ do not affect the calculated intensity.

The fractions of vertically correlated bilayers within populations 1 and 2 are denoted by *y*_1_ and *y*_2_, respectively. Thus, for population *α*, the fraction *y*_*α*_ contributes through the vertically correlated SASBIN intensity, whereas the complementary fraction 1 *− y*_*α*_ contributes through the uncorrelated bilayer intensity.

The average radius of a POPG-rich domain is not an independent fitting parameter but is calculated from the refined lattice parameter by assuming conservation of the nominal POPG molar fraction. The resulting expression is

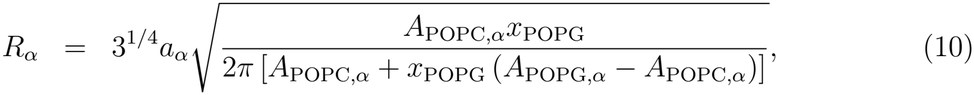

where

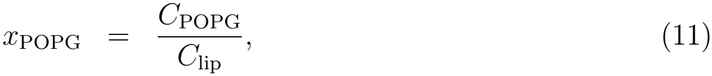

is the POPG molar fraction,

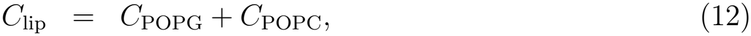

is the total lipid concentration. In the present work, *x*_POPG_ = 0.5. The concentration of membrane-associated cytochrome c in population 2 is estimated from the average area occupied by a POPG-rich domain according to

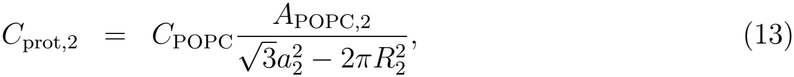

and the corresponding fraction of membrane-associated protein is

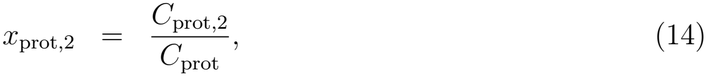

where *C*_prot_ is the nominal cytochrome c concentration approximately 0.83 mM for a molecular mass of 12 kDa, corresponding to 10 mg mL^*−*1^).

The macroscopic differential scattering cross section is described as

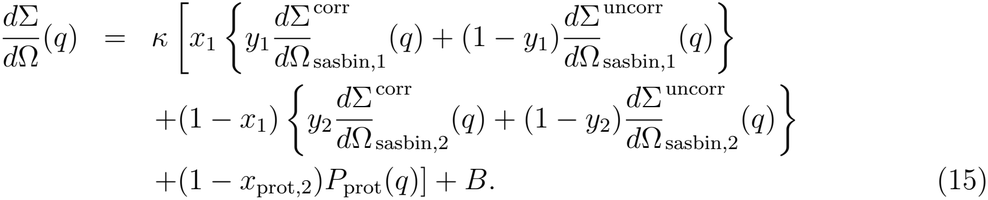

Here *κ* and *B* are a scaling factor and a flat background, respectively, accounting for instrumental effects. The quantities 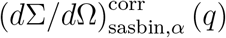 and 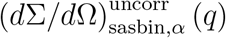 denote the scattering functions of population *α*, calculated with *N*_*c,α*_ *>* 1 and *N*_*c,α*_ = 1, respectively, according to the SASBIN model (Eq. 52 of Ref. [17]). The term *P*_prot_(*q*) is the orientationally averaged form factor of cytochrome c and accounts for the scattering contribution of protein molecules remaining free in solution. The parameter *x*_1_ is the fraction of bilayers belonging to the protein-free population, whereas 1 *− x*_1_ is the fraction of bilayers belonging to the cytochrome-c-associated population. The parameters *y*_1_ and *y*_2_ describe the degree of vertical organization within each population and are therefore distinct from *x*_1_, which describes the redistribution between the two bilayer populations.

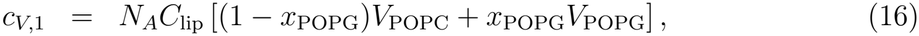

for population 1 and

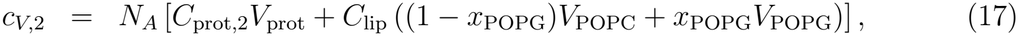

for population 2. Here *V*_POPC_ and *V*_POPG_ are the molecular volumes of the anhydrous POPC and POPG molecules, respectively, calculated as the sum of the volumes of the chemical groups listed in Table 1. The quantity *V*_prot_ is the molecular volume of cytochrome c, and *N*_*A*_ is Avogadro’s number. The last term in Eq. 15 accounts for the scattering contribution of cytochrome c molecules remaining free in solution.

All SAXS profiles, including the scattering curve recorded before the addition of cytochrome c, were analyzed simultaneously by means of a global refinement using GEN-FIT. The profile collected before mixing was described exclusively in terms of population 1, whereas all subsequent profiles were fitted according to Eq. 15. Since the SAXS curves were recorded sequentially during the interaction between cytochrome c and POPC:POPG vesicles, consecutive profiles represent nearby stages of the same structural evolution process. Accordingly, the structural parameters describing the two bilayer populations are expected to vary smoothly with time. Abrupt point-to-point fluctuations would therefore most likely reflect experimental noise or parameter correlations rather than genuine structural changes. To incorporate this physical constraint into the analysis, the time-dependent SAXS profiles were fitted simultaneously using the regularization algorithm implemented in GENFIT [20, 18]. The fitting parameters were determined by minimizing the merit function

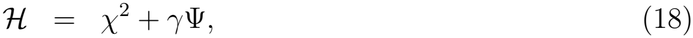

where *χ*^2^ is the average chi-square of the whole SAXS series,

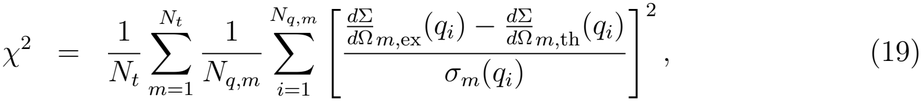

where *N*_*t*_ is the number of SAXS profiles in the time series that are fitted simultaneously, *N*_*q,m*_ is the number of experimental points of the *m*-th SAXS profile, and *σ*_*m*_(*q*_*i*_) is the corresponding experimental uncertainty of the *i*-th experimental point. The regularization functional is

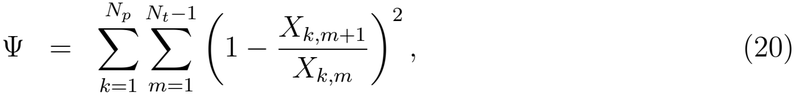

where *X*_*k,m*_ denotes the value of the *k*-th time-dependent model parameter associated with the *m*-th post-mixing SAXS profile, *N*_*p*_ is the number of time-dependent parameters included in the regularization, and *N*_*t*_ is the number of post-mixing profiles in the time series. All regularized parameters are positive over the explored range, so that the use of relative variations is well defined. The regularization term penalizes relative changes between consecutive values of the same parameter, thereby enforcing smooth temporal trajectories without imposing a common absolute scale on parameters with different units and magnitudes. The regularization was applied only to the parameters treated as time-dependent in the final refinement, whereas the scale factor *κ*, the flat background *B*, and the global time-independent parameters were excluded from the regularization term. The weighting factor *γ* was selected so that, for values of *χ*^2^ close to unity, the contribution of the regularization term remained below approximately 10% of the total merit function *H*. Under these conditions, the regularization suppresses spurious oscillations of the time-dependent parameters while preserving the quality of the fit to the experimental SAXS profiles. Preliminary refinements were used to identify parameters that were weakly determined on a curve-by-curve basis or that displayed non-systematic oscillations despite temporal regularization. In the final refinement, these parameters were treated as time-independent parameters and optimized within the global fit. Parameters describing the post-mixing profiles were constrained to be common to all SAXS curves collected after cytochrome c addition, whereas the parameters carrying the subscript “bm” were used only for the reference profile collected before mixing. Only the parameters required to describe the time-dependent redistribution between bilayer populations, protein association, lateral organization of POPG-rich domains, and vertical correlations were allowed to vary with time and were subjected to temporal regularization.

## 3. Results and Discussions

Figure 3 shows the simultaneous fit of the SAXS profiles collected before mixing and at different incubation times after the addition of cytochrome c to POPC:POPG large unilamellar vesicles. The model reproduces both the broad bilayer contribution observed before protein addition (commonly associated with bilayer form-factor scattering), and the progressive development of the lamellar stacking features appearing after mixing. In particular, the calculated curves follow the position, width, and relative intensity of the correlation peaks over the whole time series, while retaining a consistent description of the underlying bilayer form factor. This agreement indicates that the adopted two-population model captures the main structural changes induced by cytochrome c binding.

**Figure 3:**
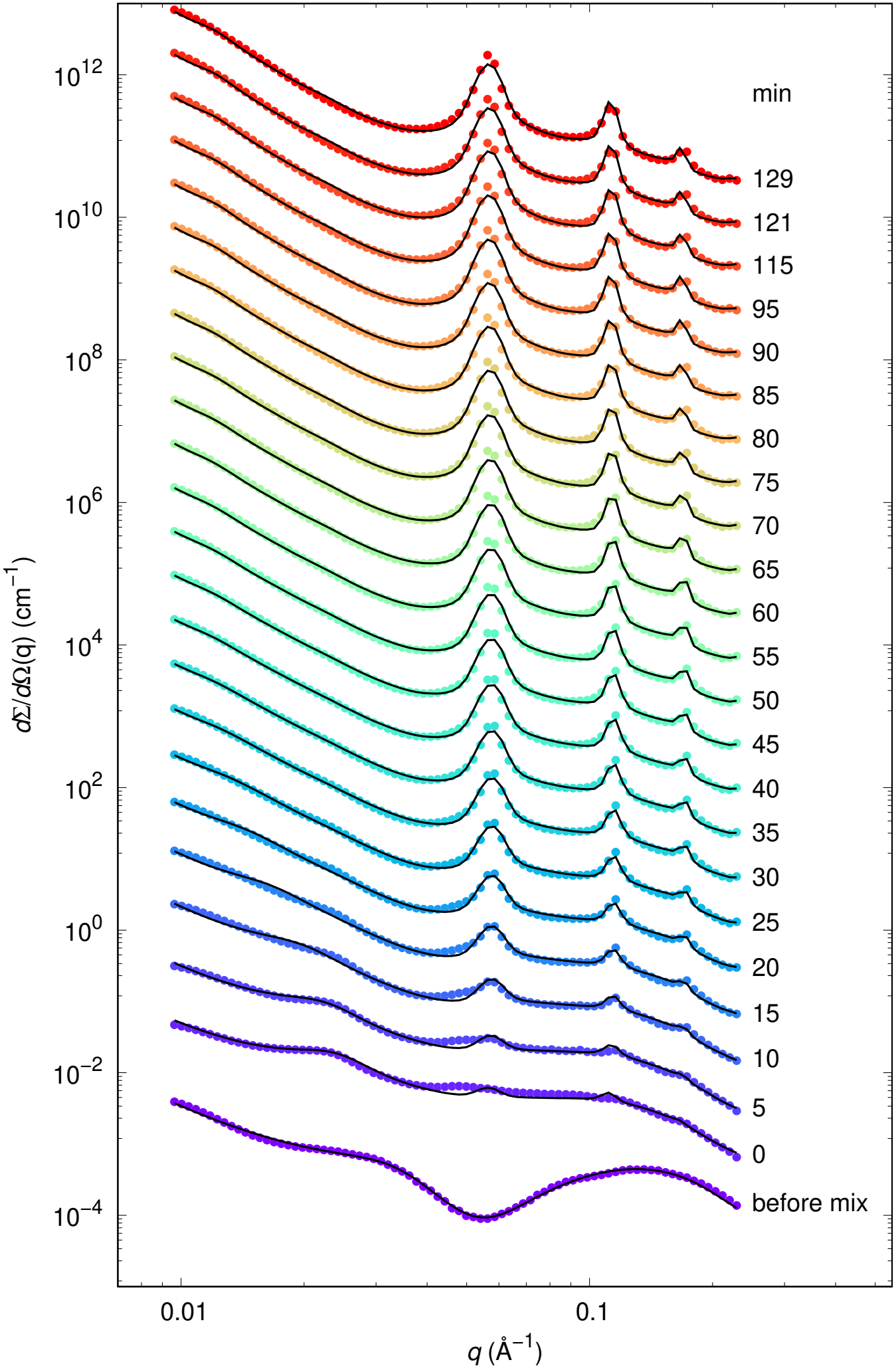
Experimental SAXS profiles collected before mixing and at different after the addition of cytochrome c to POPC:POPG large unilamellar vesicles. Open circles are experimental data and solid lines are the best global fit obtained using the SASBIN model. Curves are vertically shifted for clarity. The experimental uncertainties are smaller than the experimental points size.

A key aspect of the final refinement is that not all model parameters were allowed to vary independently along the time series. Preliminary fits showed that several molecular-profile and structural parameters were either weakly determined on a curve-by-curve basis or displayed non-systematic oscillations despite temporal regularization. These parameters were therefore constrained to be common to the relevant set of SAXS curves while still being optimized in the global fit. The remaining time-dependent parameters were those required to describe the redistribution between bilayer populations, the association of cytochrome c, the lateral organization of the POPG-rich domains, and the evolution of vertical correlations.

The time-independent parameters obtained from the final refinement are reported in Table 2. These parameters were optimized together with the time-dependent ones, but were constrained either to be common to all post-mixing SAXS curves or, for the parameters carrying the subscript “bm”, to describe only the before-mixing reference curve. Their role is there-fore to define the molecularly constrained electron-density profiles and the time-independent structural framework used in the global analysis, rather than to describe separate kinetic processes.

**Table 2:** Time-independent parameters optimized in the final refinement. Molecular group volumes were optimized globally and used to construct the constrained electron-density profiles. The parameters listed as shared by the post-mixing SAXS curves were constrained to be identical for all profiles collected after cytochrome *c* addition. Parameters carrying the subscript “bm” were instead used only to describe the reference SAXS curve collected before mixing and were treated separately within the global fit. Units are given in parentheses after the parameter symbols; parameters without units are dimensionless.

| Parameter | Value | Parameter | Value |
|---|---|---|---|
| *Molecular group volumes optimized globally* |  |  |  |
| ν<sub>CH</sub> (Å<sup>3</sup>) | 20 ± 2 | ν<sub>CH<sub>2</sub></sub> (Å<sup>3</sup>) | 25 ± 3 |
| ν<sub>CH<sub>3</sub></sub> (Å<sup>3</sup>) | 49 ± 6 | ν<sub>CG</sub> (Å<sup>3</sup>) | 139.0 ± 0.9 |
| ν<sub>CholCH<sub>3</sub></sub> (Å<sup>3</sup>) | 95 ± 2 | ν<sub>PCN</sub> (Å<sup>3</sup>) | 85.7 ± 0.8 |
| ν<sub>PGH</sub> (Å<sup>3</sup>) | 141 ± 1 |  |  |
| *Time-independent parameters shared by the post-mixing SAXS curves* |  |  |  |
| *A*<sub>POPC,1</sub> (Å<sup>2</sup>) | 67 ± 1 | *A*<sub>POPC,2</sub> (Å<sup>2</sup>) | 52.3 ± 0.3 |
| *A*<sub>POPG,1</sub> (Å<sup>2</sup>) | 80 ± 10 | *a*<sub>1</sub> (Å) | 11 ± 2 |
| *R*<sub>1</sub> (Å) | 4.1 ± 0.9 |  |  |
| *g*<sub>*a*,1</sub> | 0.05 ± 0.02 | *N*<sub>*c*,1</sub> | 16 ± 2 |
| *N*<sub>*c*,2</sub> | 5.0 ± 0.2 | *y*<sub>1</sub> | 1.0 ± 0.1 |
| *n*<sub>*w*,POPC,1</sub> | 30 ± 4 | *n*<sub>*w*,POPC,2</sub> | 22.6 ± 0.8 |
| *n*<sub>*w*,POPG,1</sub> | 4.4 ± 0.2 | *n*<sub>*w*,POPG,2</sub> | 23 ± 2 |
| *Parameters associated with the before-mixing reference curve* |  |  |  |
| *c*<sub>1,bm</sub> (Å) | 146 ± 3 | *g*<sub>*c*,1,bm</sub> | 0.41 ± 0.01 |
| *N*<sub>*c*,1,bm</sub> | 10 ± 1 |  |  |
| *Hydrophobic-core partition parameters* |  |  |  |
| φ<sub>2,POPC,1</sub> | 0.87 ± 0.02 | φ<sub>3,POPC,1</sub> | 0.20 ± 0.09 |
| φ<sub>2,POPC,2</sub> | 0.722 ± 0.009 | φ<sub>3,POPC,2</sub> | 0.40 ± 0.05 |
| φ<sub>2,POPG,1</sub> | 0.78 ± 0.04 | φ<sub>3,POPG,1</sub> | 0.27 ± 0.07 |
| φ<sub>2,POPG,2</sub> | 0.70 ± 0.08 | φ<sub>3,POPG,2</sub> | 0.400 ± 0.001 |

**Legend.** ν<sub>*j*</sub>: molecular volume of group *j* (CH, methine; CH<sub>2</sub>, methylene; CH<sub>3</sub>, terminal methyl; CG, carbonyl–glycerol; CholCH<sub>3</sub>, choline with three methyl groups; PCN, phosphate–ethylene–nitrogen; PGH, phosphoglycerol). α = 1, 2: protein-free and protein-associated bilayer populations, respectively. ℓ: lipid species (POPC or POPG). *A*<sub>ℓ,α</sub>: molecular area per lipid. *a*<sub>1</sub>: lateral hexagonal lattice parameter. *R*<sub>1</sub>: mean POPG-rich domain radius. *g*<sub>*a*,1</sub>: relative lateral paracrystalline distortion. *N*<sub>*c*,α</sub>: mean number of vertically correlated bilayers. *y*<sub>1</sub>: fraction of vertically correlated bilayers within population 1. *n*<sub>*w*,ℓ,α</sub>: number of hydration water molecules per lipid headgroup. *c*<sub>1,bm</sub>: bilayer midplane spacing before mixing. *g*<sub>*c*,1,bm</sub>: relative vertical paracrystalline distortion before mixing. *N*<sub>*c*,1,bm</sub>: mean number of vertically correlated bilayers before mixing. φ<sub>2,ℓ,α</sub> and φ<sub>3,ℓ,α</sub>: fractions of CH/CH<sub>2</sub> and terminal CH<sub>3</sub> groups, respectively, assigned to hydrophobic density level 2. bm: before mixing. Units are given in parentheses; the remaining parameters are dimensionless.

The optimized molecular group volumes are used in the molecular constraints of the model, described above in Eqs. 1-8. In these equations, the thicknesses and electron densities of the polar headgroup, hydrocarbon-chain, and terminal-methyl regions are not refined independently, but are calculated from group volumes, electron counts, molecular areas, and hydration numbers. The values reported in Table 2 therefore define the molecular building blocks of the constrained electron-density profiles used in the global fit. This strategy follows the general rationale of self-consistent bilayer models, in which area per lipid, hydration, bilayer thickness, and electron density are linked by volume conservation rather than treated as unrelated fitting parameters [13, 14, 10, 11, 15]. The optimized hydrocarbon group volumes are close to the values commonly adopted in SDP-type descriptions of phospholipid bilayers. In particular, *ν*_CH_ = 20 *±* 2 Å^3^ is essentially unchanged with respect to the reference value, whereas *ν* CH_2_ = 25 *±* 3 Å^3^ and 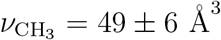 are slightly smaller but remain within the range expected for effective molecular volumes in a constrained global fit.

The refined molecular areas per lipid headgroup provide an internal check on the bilayer profiles associated with the two populations. The area of the POPC-rich matrix in population 1, *A*_POPC,1_ = 67*±*1 Å^2^, is close to values commonly reported for fluid phosphatidylcholine bilayers with unsaturated chains [10, 11, 12]. This agreement supports the interpretation of population 1 as a protein-free POPC-rich membrane with fluid-like molecular packing.

The area associated with POPG in the same population, *A*_POPG,1_ = 80 *±* 10 Å^2^, is larger; however, this value should not be interpreted as the area of an extended POPG-rich phase. Indeed, the corresponding POPG-rich radius, *R*_1_ = 4.1 *±* 0.9 Å, is comparable with a molecular length scale. Together with the small lateral lattice parameter *a*_1_ = 11 *±* 2 Å, this is consistent, within the adopted model, with the absence of resolved extended POPG-rich domains. It does not constitute a direct measurement of molecular lipid dispersion.

In population 2, the POPC molecular area, *A*_POPC,2_ = 52.3 *±* 0.3 Å ^2^, is markedly smaller than in population 1. This value should be interpreted as an effective molecular area of the POPC-rich matrix in the cytochrome-c-associated bilayers, rather than as the area of a pure POPC membrane. Its reduction is consistent with a more compact interfacial environment, where electrostatic binding, POPG-rich domains, headgroup hydration, and bilayer packing are coupled within the constrained electron-density profile.

Hydration numbers are effective parameters inferred jointly with molecular areas and electron-density profiles, rather than independent measurements of interfacial water. Their interpretation is conditional on the assumed molecular volumes, population assignment, and parameter correlations.

Within the assumed electron-density description, the hydration parameters suggest that the two populations correspond to distinct interfacial environments. The POPC-rich matrix in population 1 is highly hydrated, *n*_*w*,POPC,1_ = 30 *±* 4, whereas POPC in population 2 has a lower but still substantial hydration, *n*_*w*,POPC,2_ = 22.6 *±* 0.8. This decrease parallels the smaller effective POPC area in population 2 and is consistent with a more compact protein-associated interfacial environment.

The contrast is even more pronounced for POPG. In population 2, *n*_*w*,POPG,2_ = 23 *±* 2, showing that the POPG-rich domains associated with cytochrome c remain substantially hydrated. This supports a picture in which the protein is accommodated at a hydrated membrane interface rather than forming a completely dehydrated contact with the anionic domains. By contrast, the much smaller value obtained for population 1, *n*_*w*,POPG,1_ = 4.4 *±* 0.2, should not be interpreted as the hydration of an extended POPG-rich phase. In this population, the POPG contribution is reduced to a molecular-scale effective inhomogeneity, and the hydration number mainly controls the electron-density contrast of dispersed anionic lipids within the POPC-rich matrix.

The remaining global parameters define structural features that were not allowed to vary during the post-mixing time series. The small values of *a*_1_ = 11 *±* 2 Å and *R*_1_ = 4.1 *±* 0.9 Å, together with the low lateral distortion parameter, *g*_*a*,1_ = 0.05 *±* 0.02, support the interpretation of population 1 as a protein-free membrane in which POPG is effectively dispersed within the POPC-rich matrix rather than organized into extended anionic domains. The average numbers of correlated bilayers, *N*_*c*,1_ and *N*_*c*,2_, 16 *±* 2 and 5.0 *±* 0.2, were also treated as global parameters. Therefore, changes in vertical organization after cytochrome c addition are described primarily by the time-dependent fractions, correlation distances, and paracrys-talline distortion parameters, rather than by a changing average stack size. The parameters carrying the subscript “bm” describe the SAXS curve collected before cytochrome c addition.

They were optimized within the global refinement but were used only for the before-mixing reference profile. The values *c*_1,bm_ = 146 *±* 3 Å, *g*_*c*,1,bm_ = 0.41 *±* 0.01, and *N*_*c*,1,bm_ = 10 *±* 1 therefore define the initial lamellar organization of the lipid dispersion before protein addition and are not constrained to be shared with the post-mixing profiles. The hydrophobic-core partition parameters *ϕ*_2_ and *ϕ*_3_ define how the hydrocarbon-chain and terminal-methyl contributions are assigned to the two hydrophobic electron-density levels of the model. They are therefore part of the molecular definition of the bilayer profile. The refined values remain far from the limiting case *ϕ*_2_ = 1 and *ϕ*_3_ = 0, and therefore do not describe a sharply segregated hydrophobic core. Rather, they indicate a partial overlap between chain and terminal-methyl contributions, as expected for fluid bilayers with conformationally disordered acyl chains.

The trajectories below describe the evolution within the adopted model. Their smoothness is partly imposed by temporal regularization and does not itself establish distinct rate constants or a sequential mechanism. Protein-associated fractions must be interpreted using the sample-wide mass balance discussed above.

The time-dependent parameters retained in the final regularized refinement are shown in Fig. 4. These parameters describe the time-dependent component of the model. The population fraction *x*_1_ decreases rapidly after cytochrome c addition, indicating a progressive redistribution of bilayers from the protein-free population 1 to the cytochrome-c-associated population 2. This redistribution occurs mainly during the first 30-40 min and then approaches a plateau. In parallel, the model-derived fraction of membrane-associated protein, *x*_prot,2_, increases rapidly and reaches an approximately constant value on a similar time scale. The simultaneous decrease of *x*_1_ and increase of *x*_prot,2_ indicate that the formation of population 2 is coupled to cytochrome c association rather than representing a purely lipid-only structural rearrangement.

**Figure 4:**
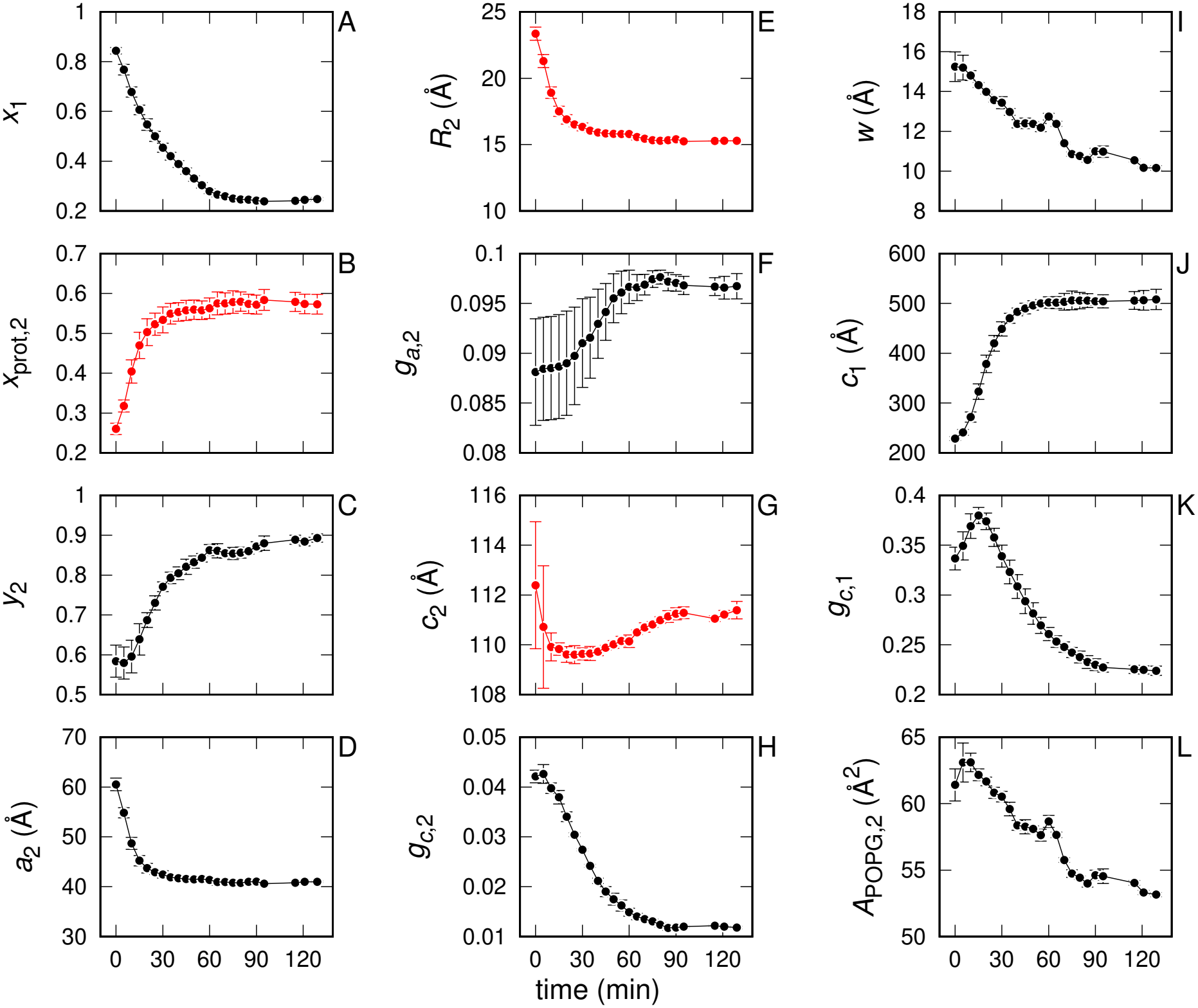
Time evolution of the parameters allowed to vary along the SAXS time series in the final regularized refinement. Panels report: (A) fraction *x*_1_ of bilayers belonging to population 1; (B) model-derived fraction *x*_prot,2_ of membrane-associated cytochrome c in population 2; (C) fraction *y*_2_ of vertically correlated bilayers in population 2; (D) lateral lattice parameter *a*_2_ of the POPG-rich domains in population 2; (E) corresponding POPG-rich domain radius *R*_2_; (F) lateral paracrystalline distortion parameter *g*_*a*,2_; (G) vertical correlation distance *c*_2_ of population 2; (H) vertical paracrystalline distortion parameter *g*_*c*,2_ of population 2; (I) water thickness *w* separating cytochrome c from the POPG-rich domain; (J) vertical correlation distance *c*_1_ of population 1; (K) vertical paracrystalline distortion parameter *g*_*c*,1_ of population 1; and (L) molecular area *A*_POPG,2_ of POPG in population 2. Red symbols denote quantities calculated from the fitted parameters, whereas black symbols denote directly refined parameters. Error bars are standard uncertainties from the fitting procedure or propagated uncertainties for derived quantities.

The fraction *y*_2_ of vertically correlated bilayers within population 2 also increases with time. Within the model, population redistribution and vertical correlations are described by separate parameters; their differing trajectories suggest coupled structural changes but do not independently establish distinct kinetic processes. The former is described by the redistribution between populations, through *x*_1_, whereas the latter is described by the increasing weight of the correlated component inside population 2, through *y*_2_. Thus, cytochrome c binding is associated not only with the appearance of a structurally distinct membrane population, but also with a progressive increase in the fraction of that population participating in vertically correlated stacks.

The lateral organization of the POPG-rich domains in population 2 undergoes a pro-nounced early change. The hexagonal lattice parameter *a*_2_ decreases sharply during the first part of the experiment and then levels off. The corresponding domain radius *R*_2_, calculated from the conservation of the nominal POPG molar fraction, follows the same trend. Since *R*_2_ is a derived quantity, its time dependence should be interpreted together with that of *a*_2_ and of the molecular area *A*_POPG,2_. Nevertheless, the decrease of both *a*_2_ and *R*_2_ indicates that, in the protein-associated population, the POPG-rich domains become more compactly organized after cytochrome c binding. Within the adopted model, this behavior is consistent with a cytochrome-c-associated reorganization of the anionic lipid-rich regions, although it does not distinguish the formation of new domains from the remodeling of pre-existing heterogeneity.

The lateral paracrystalline distortion parameter *g*_*a*,2_ shows a more limited but systematic evolution. After an initial regime with larger uncertainty, *g*_*a*,2_ increases slightly and then remains nearly constant. This suggests that, once the characteristic lateral spacing of the POPG-rich domains has decreased, their degree of lateral disorder reaches a quasi-stationary value. Therefore, the dominant lateral effect is the reduction of the domain spacing and radius, whereas the change in lateral disorder is secondary.

The vertical organization of population 2 evolves differently from the lateral one. The correlation distance *c*_2_ varies only moderately over the time series, remaining close to the value imposed by the geometry of the protein-mediated stack. In contrast, the vertical paracrystalline distortion parameter *g*_*c*,2_ decreases markedly, especially during the first part of the experiment. This decrease indicates that the cytochrome-c-associated stacks become progressively better ordered along the bilayer normal. At the same time, the water thickness *w* between the protein and the POPG-rich domain decreases, suggesting that the protein-associated bilayers undergo a gradual compaction of the interlamellar region. Taken together, the trends of *y*_2_, *g*_*c*,2_, *c*_2_, and *w* indicate that vertical ordering develops more gradually than the initial redistribution between the two bilayer populations.

Although population 1 becomes progressively less abundant after cytochrome c addition, its vertical parameters still provide useful information on the residual protein-free bilayers. The correlation distance *c*_1_ increases substantially with time, whereas *g*_*c*,1_ initially increases and then decreases toward a lower plateau. Because the weight of population 1 decreases strongly during the same time interval, these parameters should be interpreted with caution. Likewise, a fitted radius on the order of 4 Å represents a molecular-scale effective contrast parameter; it does not directly resolve individual POPG molecules or prove molecular dispersion. They mainly indicate that the residual protein-free bilayers are not simply static spectators, but are affected by the global reorganization of the sample as cytochrome c redistributes the membrane material into the protein-associated population.

The only molecular-profile parameter retained as time-dependent in the final analysis is *A*_POPG,2_. Its decrease indicates that the effective molecular profile of POPG in the protein-associated domains changes during the same time interval in which the lateral domain radius and the protein–membrane separation decrease. This parameter should not be overinterpreted as an isolated measurement of lipid area, because it is coupled to the molecular-volume constraints and to the calculation of the domain radius. Rather, it reports that the electron-density profile of the POPG-rich domains in population 2 must change in order to reproduce the time evolution of the SAXS curves. The decrease of *A*_POPG,2_ also contrasts with the larger effective value obtained for *A*_POPG,1_ in the protein-free population. This contrast reinforces the view that POPG plays different structural roles in the two populations: in population 1 it behaves as an effectively dispersed contribution within the POPC-rich matrix, whereas in population 2 it forms more compact POPG-rich domains associated with cytochrome c.

Overall, the fitted trajectories suggest early lateral reorganization and population redistribution, together with a more gradual evolution of interbilayer correlations. These observations are consistent with a coupling between cytochrome c association, lateral POPG-rich organization, and lamellar ordering within the adopted two-population model. Although this model consistently describes the experimental SAXS profiles, the present analysis does not establish its uniqueness without a systematic comparison with alternative structural descriptions. Moreover, the data do not distinguish between POPG-rich domain formation and the stabilization of pre-existing heterogeneity.

The structural evolution observed in this study provides new insights into the interplay between cytochrome c association, anionic lipid organization, and membrane remodeling, processes relevant to mitochondrial function and apoptotic signaling. Recent studies have demonstrated that cytochrome c interactions with negatively charged phospholipids extend beyond electrostatic adsorption, involving specific binding configurations, protein conformational changes, and membrane structural rearrangements. Elmer-Dixon and Bowler [21] highlighted the importance of electrostatic interactions at the protein–membrane interface, whereas Mohammadyani et al. [22] investigated cardiolipin-induced structural perturbations associated with cytochrome c binding. Complementary spectroscopic investigations by Ripanti et al. [23] and Sun et al. [24] further demonstrated the complexity of cardiolipin-mediated interactions, emphasizing their influence on protein conformation and membrane association. More recently, Chow et al. [25] used hydrogen–deuterium exchange mass spectrometry to characterize cytochrome c interactions with anionic phospholipid nanodiscs, providing additional insights into membrane-binding regions. Furthermore, Li et al. [26] demonstrated that different anionic phospholipids can induce distinct conformational and functional responses in cytochrome c. In this context, our time-resolved SAXS results offer a complementary perspective by examining the structural response of lipid assemblies rather than protein conformational changes. Within the molecularly constrained SASBIN framework, the scattering profiles are consistently described by two bilayer populations, with protein association accompanied by changes in lateral POPG-rich organization and progressively enhanced lamellar ordering. The fitted trajectories suggest that lateral reorganization and population redistribution are particularly pronounced during the initial stages, whereas interbilayer correlations evolve more gradually. These findings support the view that anionic membranes are not passive binding surfaces but dynamically reorganizing interfaces whose structural properties are coupled to peripheral protein association. Although POPC:POPG membranes do not reproduce the chemical complexity of mitochondrial membranes, our results provide a framework for investigating how protein binding influences collective membrane organization and its potential relationship with lipid-dependent biological functions.

## 4. Conclusions

Time-resolved SAXS interpreted with a molecularly constrained two-population SAS-BIN model provides a structural description of the response of POPC:POPG dispersions to cytochrome c addition. Within this framework, protein-associated bilayers exhibit effective lateral correlations consistent with POPG-rich regions and an evolution toward greater lamellar ordering. The molecular-scale inhomogeneities assigned to the protein-free population should be treated as effective scattering contributions rather than directly resolved domains. The fitted trajectories suggest that lateral reorganization and population redistribution are prominent during the first tens of minutes, while some vertical correlation parameters evolve more gradually. These trends support coupling between membrane association and structural remodeling, but do not establish independent kinetic pathways or a unique temporal sequence. The necessity of two populations, quantitative protein association, and robustness of the inferred structural changes remain conditional on model comparison, consistent protein conservation, and sensitivity analysis. The broader implication is that peripheral protein association can be examined in terms of coupled lateral and interbilayer organization. Independent morphology and binding measurements would strengthen this interpretation. Extrapolation from this POPC:POPG system to cardiolipin-containing mitochondrial membranes or apoptotic function requires additional evidence. More generally, the molecularly constrained SASBIN framework may be extended to other membrane systems exhibiting lateral heterogeneity and interbilayer correlations, and could potentially be adapted to small-angle neutron scattering through an appropriate description of the scattering-length-density profiles.

## 5. Acknowledgements

This work was supported by grants from the São Paulo Research Foundation (FAPESP, grant numbers 2022/07231-7 e 2023/04532-9), the Brazilian National Council for Scientific and Technological Development (CNPq, grant numbers 401982/2025-4, 309359/2025-2, 311831/2021-4). This research used the SAXS beamline of the Brazilian Synchrotron Light Laboratory (LNLS), part of the Brazilian Center for Research in Energy and Materials (CN-PEM), Campinas, Brazil, we also thank the SAXS beamline staff for technical support. The authors declare the use of AI-assisted tools during the preparation of this manuscript. All content, including all bibliographic analysis, was critically reviewed, verified, and approved by the authors, who take full responsibility for the accuracy and integrity of the work.

